# Use antibiotics in cell culture with caution: genome-wide identification of antibiotic-induced changes in gene expression and regulation

**DOI:** 10.1101/130484

**Authors:** Ann H. Ryu, Walter L. Eckalbar, Anat Kreimer, Nir Yosef, Nadav Ahituv

**Affiliations:** Department of Bioengineering and Therapeutic Sciences, University of California San Francisco, San Francisco, CA, USA; Institute for Human Genetics, University of California San Francisco, San Francisco, CA, USA; Department of Electrical Engineering and Computer Science and the Center for Computational Biology, University of California Berkeley, Berkeley, CA, USA; Ragon Institute of Massachusetts General Hospital, Massachusetts Institute of Technology, and Harvard University, 02139 Boston, Massachusetts, USA

## Abstract

Standard cell culture guidelines often use media supplemented with antibiotics to prevent cell contamination. However, relatively little is known about the effect of antibiotic use in cell culture on gene expression and the extent to which this treatment could confound results. To comprehensively characterize the effect of antibiotic treatment on gene expression, we performed RNA-seq and ChIP-seq for H3K27ac on HepG2 cells, a human liver cell line commonly used for pharmacokinetic, metabolism and genomic studies, cultured in media supplemented with penicillin-streptomycin (PenStrep) or vehicle control. We identified 205 PenStrep-responsive genes, including transcription factors such as *ATF3* that are likely to alter the regulation of other genes. Pathway analyses found a significant enrichment for “xenobiotic metabolism signaling” and “PXR/RXR activation” pathways. Our H3K27ac ChIP-seq identified 9,514 peaks that are PenStrep responsive. These peaks were enriched near genes that function in cell differentiation, tRNA modification, nuclease activity and protein dephosphorylation. Our results suggest that PenStrep treatment can significantly alter gene expression and regulation in a common liver cell type such as HepG2, advocating that antibiotic treatment should be taken into account when carrying out genetic, genomic or other biological assays in cultured cells.

## Introduction

One of the most common cautionary measures taken during *in vitro* studies is the use of antibiotics while culturing cells in order to avoid bacterial contamination. Standard cell culture protocols listed by the American Type Culture Collection (ATCC) explicitly entail the addition of antibiotics, such as penicillin-streptomicin (PenStrep) and gentamicin, as media supplements ^1^. Many large-scale genomic projects, such as the ENCODE project ^2^, use cell lines in order to understand the diversity of gene expression and regulatory profiles across human cell types and require the routine use of antibiotics in their protocols. The implicit assumption made within the community is that using antibiotics in cell culture has a negligible impact on gene expression.

Previous studies have demonstrated that changes in gene expression and regulation *in vitro* can be induced by antibiotics ^3,4^. One study has even shown that antibiotics such as rifampin can induce genome-wide, drug-dependent changes in gene regulation and expression patterns in human hepatocytes. However, the molecular consequences of growing human cells with antibiotics at standard cell culture concentrations have yet to be thoroughly investigated.

Penicillin is a group of antibiotics metabolized in the liver that eliminate bacteria by inhibiting the peptidoglycan synthesis necessary to maintain the bacterial cell wall ^5^. Streptomycin is an antibiotic that acts as a protein synthesis inhibitor by binding to the small 16S rRNA of the 30S subunit of the bacterial ribosome, interfering with codon reading and ultimately the death of microbial cells through mechanisms that are still not well understood ^6^. Although streptomycin is used in combination with penicillin in standard antibiotic cocktails to prevent bacterial infection in cell culture, their mechanism of action in cells other than microbial cells are not well understood. In order to investigate the effects of antibiotics commonly used in cell culture on gene expression and regulation, we performed RNA-seq and H3K27ac ChIP-seq on HepG2 cells, immortalized human liver cells, treated with or without PenStrep. We identified a set of differentially expressed genes responsive to PenStrep that was significantly enriched for pathways involved in drug metabolism and known pathways induced by gentamicin, another commonly used antibiotic in cell culture. ChIP-seq for H3K27ac, an active promoter and enhancer mark ^7,8^, identified thousands of PenStrep responsive peaks. Combined, our study elucidates genes and regulatory regions that are activated due to antibiotic treatment in liver cells, and suggests that the use of antibiotics in genomic and other biological studies should be taken into account in any type of analysis.

## Results

### Drug-associated genes are differentially expressed following PenStrep treatment

To systematically identify genes that are differentially expressed due to cell culture antibiotic treatment, we compared gene expression levels of HepG2 cells cultured with standard 1% PenStrep-supplemented media and HepG2 cells cultured with media not supplemented with PenStrep. Using DESeq2 to perform differential expression analysis ^9^, we identified 205 differentially expressed (DE) genes using a q-value cutoff, after adjustment for multiple testing of less than or equal to 0.1 (Fig. 2a, Table S1). Amongst these DE genes, 156 were significantly upregulated due to PenStrep treatment and 49 were downregulated (Fig.1). These include a set of transcription factors- *ATF3, SOX4, FOXO4, TGIF1, HOXD1, FOXC1, GTF3C6* – some of which are known to play a significant role in drug and stress response ^4,10–23^. Pathway analysis using the Database for Annotation, Visualization and Integrated Discovery (DAVID; ^24^) found that the cluster of 156 DE genes that were upregulated due to PenStrep are enriched for gene ontology terms associated with apoptosis (p-value = 1.91E-05), drug response (p-value = 1.58E-04), unfolded protein response (p-value = 3.84E-04), and nitrosative stress (p-value = 3.98E-04) (Fig. 2a). DE genes that were downregulated in response to PenStrep were enriched for gene ontology categories related to insulin response (p-value = 6.85E-04), cell growth and proliferation (p-value = 0.012), toxic substance (p-value = 0.018) and drug response (p-value = 0.012) (Fig. 2a). Further analyses of PenStrep DE genes using Ingenuity pathway analysis (IPA), found canonical pathway enrichment for PXR/RXR activation (p-value = 9.43E-05), a known drug response pathway associated with antibiotic treatment ^25^ (Fig. 2b). Further IPA analyses for upstream regulators enriched for DE genes, identified a significant enrichment for gentamicin (p-value = 2.93E-13), an aminoglycoside (like streptomycin) that is associated with nephrotoxicity as well as ototoxicity in human patients ^26–28^ but also commonly used in cell culture on target genes (Fig. 2b and c). The overlap between target genes of gentamicin and the PenStrep-dependent genes in our analysis demonstrates a similar mechanism of action across antibiotics in human cells. Overall, the diversity of gene pathways activated in PenStrep-treated cells not only suggests that PenStrep induces a systemic change in gene expression in human cell lines, but that PenStrep may also be inducing broader changes at the gene regulatory level.

**Figure 1.**
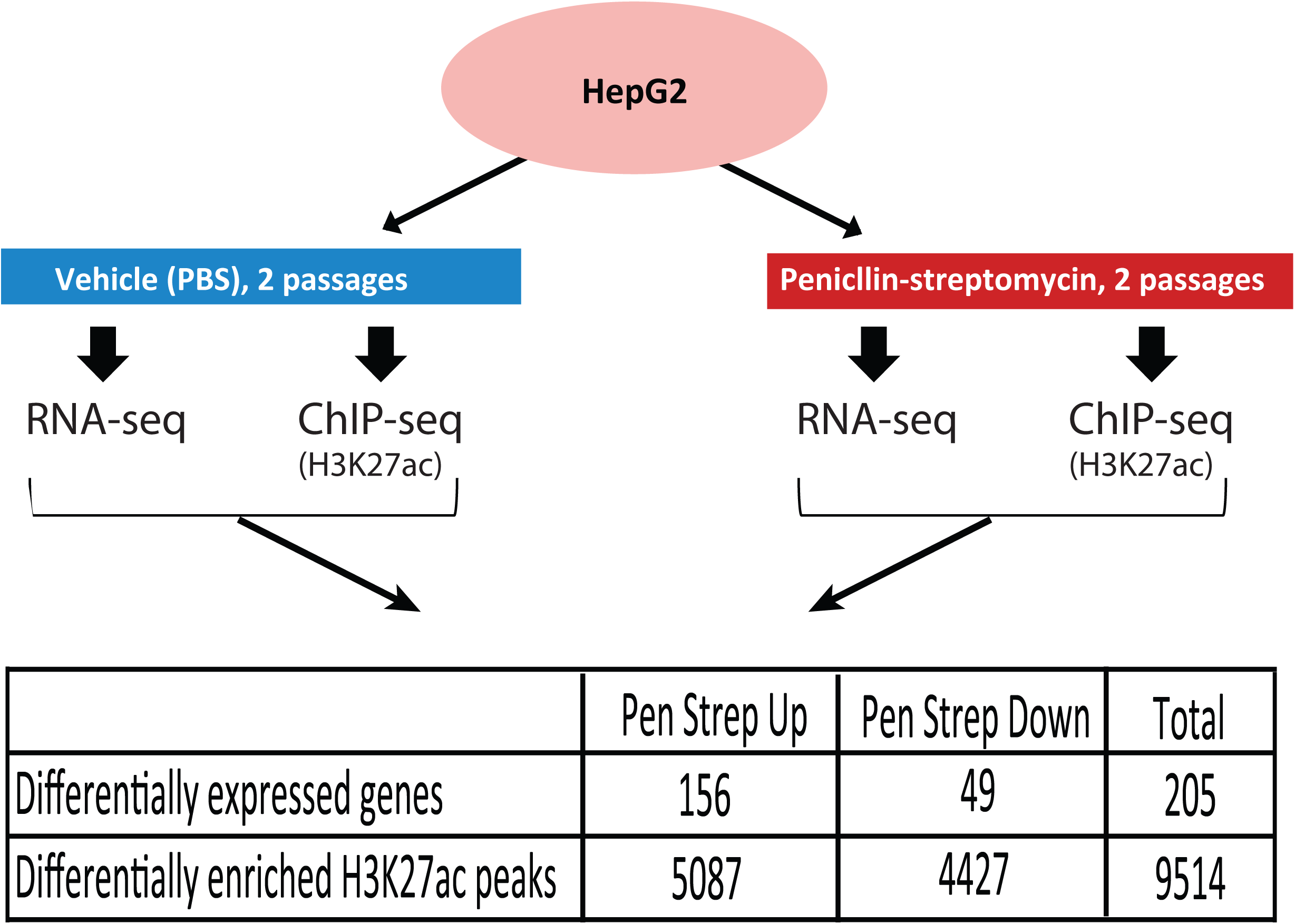
Experimental design. Schematic of the RNA-seq and ChIP-seq assays performed on HepG2 cells treated with and without PenStrep. The number of differentially expressed genes and differentially enriched H3K27ac peaks are listed in the table below the diagram.

**Figure 2.**
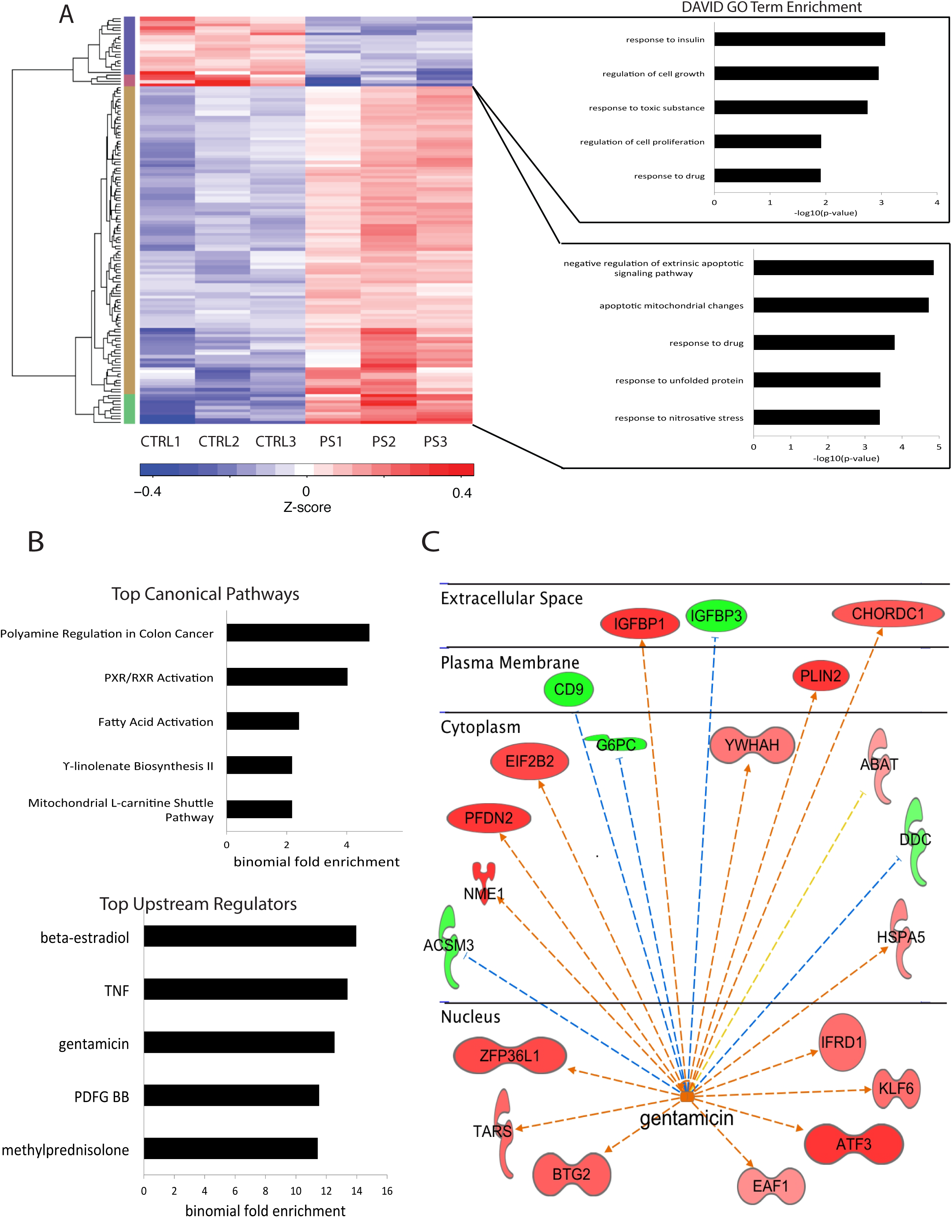
RNA-seq analysis on HepG2 cells cultured with and without PenStrep. (A) Shown in the top left is a heatmap depicting relative expression levels for all 205 differentially expressed genes across all three replicates per treatment. To the right of each cluster, the top gene ontology terms as determined by DAVID v6.8 ^24^ are shown. (B) PenStrep DE genes canonical pathways and upstream regulators with the top five highest binomial fold enrichment values and most significant p-values as determined by IPA. (C) IPA network analysis for gentamicin, one of the top most significant upstream regulators for PenStrep DE genes. This network is drawn based on calculated z-scores for published gene expression patterns under gentamicin as determined by IPA.

### PenStrep induces differential enrichment of active promoter and enhancer regions marked by H3K27ac

To determine whether culturing cells with PenStrep also leads to chromatin landscape changes that can alter gene regulation, we performed ChIP-seq for H3K27ac on both HepG2 cells cultured with and without PenStrep. Using DESeq2 ^9^, we annotated a total of 9,514 peaks that are differentially enriched between PenStrep and control treatments at a q-value cutoff, after adjustment for multiple testing, of less than or equal to 0.1. Of these peaks, 5,087 were highly enriched in the PenStrep condition and 4,427 peaks were highly enriched in the control treatment (Fig. 1 and Fig. 3a). Using the Genomic Regions Enrichment Annotation Tool (GREAT; ^29^), we identified genes nearby each cluster of DE regions separately (up or down) for gene ontology enrichment. For the cluster of enriched H3K27ac peaks induced by PenStrep, we observed a significant association with genes involved in tRNA modification (p-value= 2.0E-08), regulation of nuclease activity (p-value= 2.0E-08), cellular response to misfolded protein (p-value= 1.1E-07), and regulation of protein dephosphorylation (p-value= 1.9E-07)(Fig. 3a). As streptomycin is known to act as a protein synthesis inhibitor by binding to the small 16S rRNA of the 30S subunit of the bacterial ribosome, this suggests that the known mechanism of action for streptomycin in bacterial cells may also affect mammalian cells ^6,30^. For the cluster of H3K27ac peaks that were significantly enriched in the control treatment, GREAT identified an enrichment for genes involved in stem cell differentiation (p-value= 6.8E-22), actin depolymerization (p-value=1.3E-21), negative regulation of transcription factor activity (p-value=2.0E-19), response to reactive oxygen species and positive regulation of cell cycle (p-value=1.2E-14)(Fig. 3a). Combined, our results demonstrate a global change in the regulatory landscape induced by PenStrep that corroborate some of the gene pathways enriched in our RNA-seq results as well as toxicity pathways that are associated with streptomycin.

**Figure 3.**
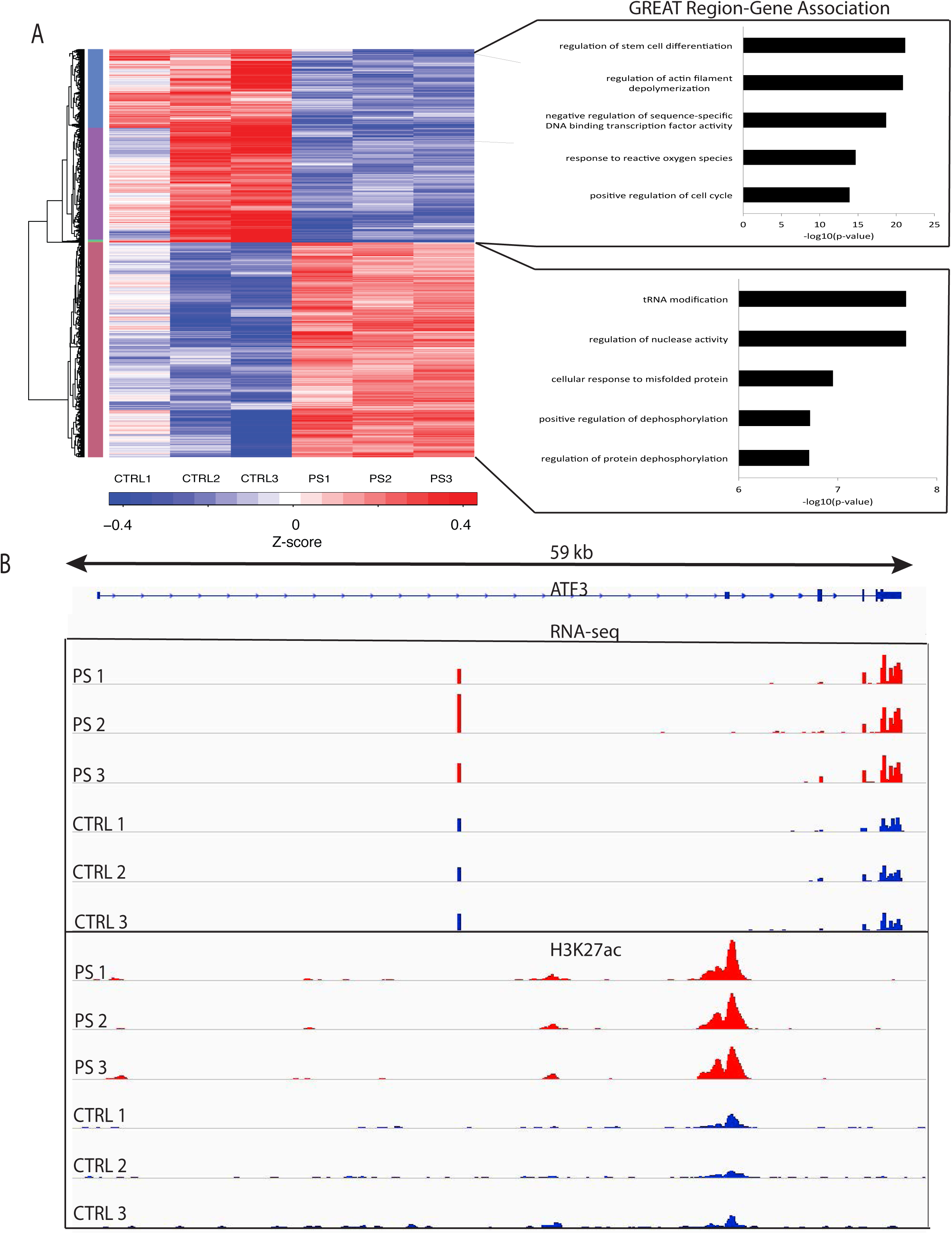
ChIP-seq analysis of PenStrep responsive peaks in HepG2 cells. (A) The top right panel shows a heatmap depicting relative expression levels for all 9,514 differentially enriched H3K27ac peaks across all three replicates per treatment. To the right, are shown the top region-gene association pathways according to GREAT ^29^. (B) Integrative genomic viewer snapshot showing RNA-seq and H3K27ac ChIP-seq results in the *ATF3* locus. This locus shows increased *ATF3* expression in the PenStrep-treated HepG2 cells (red) versus vehicle control treated cells (blue) observed through RNA-seq, as well as ChIP-based enrichment of H3K27ac in PenStrep-treated cells (red) versus vehicle control treated cells (blue).

### PenStrep-responsive regulatory regions overlap or reside near PenStrep-associated genes

In order to test for a correlation between the DE H3K27ac peaks and DE genes found via ChIP-seq and RNA-seq respectively, we took all 9,514 DE regions and matched the closest gene expressed (i.e. had a normalized FPKM greater than zero across replicates for all conditions). We found a positive Spearman correlation between regions having a DE H3K27ac signal and expression of DE genes that are enriched in the PenStrep condition (r=0.21) as well as a positive Spearman correlation between DE H3K27ac regions and DE genes that are depleted in the PenStrep condition (r=0.15). Additionally, we performed hyper geometric tests on matching clusters of PenStrep dependent DE genes and DE H3K27ac regions in order to determine significance of enrichment of DE genes within DE H3K27ac regions. We found that the cluster of DE genes and DE H3K27ac regions that are up in PenStrep have significant overlap (p-value=1.21E^-07^). Conversely, the cluster of DE genes and DE H3K27ac regions that are down in PenStrep also have significant overlap (p-value=2.75E^-08^). These findings corroborate the similar gene ontology term enrichment found for both our RNA-seq and ChIP-seq DE genes and peaks. GO terms related to misfolded protein response were found for both clusters of genes and H3K27ac peaks that showed an increase upon PenStrep treatment (Fig. 2a and Fig. 3a). Similarly, the clusters of DE genes and H3K27ac peaks that were depleted upon PenStrep treatment showed GO term enrichment for cell cycle regulation and cell growth. Many of these DE H3K27ac regions were found to overlap DE genes that are transcription factors, as is underscored by enrichment of regions near genes important for transcription factor activity found through our GREAT analysis (Fig. 3a). One notable example of this is *ATF3*, which is more highly expressed in the PenStrep condition and has a regulatory region that is also enriched in the PenStrep condition that overlaps its third exon (Fig. 3b). *ATF3* is a transcription factor that is known to play a myriad of roles in cell differentiation and proliferation ^31–34^, response to unfolded proteins ^35^, inflammation and immune response ^36–38^, and regulating hepatic gluconeogenesis and insulin resistance ^39–43^. *ATF3* is also known to be a major player in drug response ^4,10–16^. Combined, our results show correlation between DE regulatory elements and DE genes and highlight several pathways and genes that could be associated with PenStrep response.

## Discussion

Standard protocols for cell culture frequently require antibiotics to reduce contamination. Amongst the most popular antibiotics used for cell culture are gentamicin and PenStrep, a cocktail of both penicillin and streptomycin. Streptomycin and gentamicin are both aminoglycosides, a family of antibiotics that is well known to induce nephrotoxicity ^26–28^ as well as ototoxicity in patients. Despite their known toxicity in humans, there have been only a few reports that have demonstrated how these antibiotics impact gene expression. These studies have shown the impact of protein synthesis and enzyme activity in rat cell cultures ^44–46^ and that these antibiotics produce side effects during human stem cell differentiation into adipocytes ^30^. However, the systematic effects of using these antibiotics at the standard cell culture concentrations on gene expression and regulation in human cell lines have not been thoroughly investigated ^30^. Here, by carrying out RNA-seq and ChIP-seq on PenStrep and vehicle treated HepG2 cells, we show that antibiotics can induce a global change in gene expression and chromatin landscape in a human cell line.

We find that PenStrep responsive genes are not only involved in pathways related to drug response, but also to insulin response, fatty acid activation, mitochondrial l-carnitine shuttle pathways, apoptosis, cell growth, and unfolded protein response. We also observe that many of these PenStrep-dependent DE genes are also known targets of gentamicin, some of which are nuclear receptors and transcription factors (e.g. *ATF3*). Gentamicin and other members of the aminoglycoside family, such as streptomycin, are associated with both nephrotoxicity and ototoxicity in humans ^26,27^. However, the molecular mechanisms underlying death of proximal tubule kidney cells and mechanosensory hair cells due to aminoglycosides are not well understood ^28^. Given that both types of toxicity have been associated with most aminoglycosides, streptomycin could be an activator of toxicity pathways across cell types. The overlap of PenStrep-activated DE genes with known targets of gentamicin suggests that toxicity pathways known to be induced by antibiotics could exist not only in microbial cells and human patients, but also in human cells *in vitro*. Furthermore, our results may also reveal some patterns in gene expression and regulatory regions underlying toxicity pathways induced by aminoglycosides as a group.

Changes in gene expression were further corroborated by changes in regulatory regions observed in our H3K27ac ChIP-seq data from the same PenStrep treatment of HepG2 cells. Not only was there significant overlap between DE regions found in our H3K27ac data, but when we looked at genes nearby DE regions dependent on PenStrep treatment, we found enrichment for several similar pathways. Regulation of cell cycle and cellular response to misfolded protein reappeared as potential pathways regulated by H3K27ac regions activated by PenStrep. Some pathways based on H3K27ac DE region-gene associations were reminiscent of the known mechanism of action of streptomycin in bacteria, such as tRNA modification. Other pathways found to be enriched in our DE region-gene associations, such as response to reactive oxygen species and cytoskeleton depolymerization, were previously shown to be disrupted by antibiotics in the differentiation of human stem cells into adipocytes ^30^ and mouse embryonic stem cells into neurons ^47^. The recapitulation of pathways disrupted by antibiotics at standard cell culture concentrations in mouse stem cells, human stem cells, human adipocytes and the human liver cell line we used for this study, suggests that the mechanism of action of antibiotics is likely to be detectable and similarly perturbed across other human cell lines commonly used for research.

Several transcription factors were also observed to have enhanced expression due to PenStrep treatment, which could potentially alter the expression of additional genes and pathways. Some of these transcription factors are known to be key regulators of differentiation, drug response, and cell cycle and growth. One of the most notable transcription factors amongst our observed PenStrep – dependent genes, is *ATF3*, a transcriptional repressor that is known to play a role in cell differentiation and proliferation ^31–34^, the unfolded protein response ^35^, inflammation and immune response ^36–38^, hepatic gluconeogenesis and insulin resistance ^39–43^, as well as drug response ^4,10–16^. *ATF3* is well-studied in a wide diversity of human cell types including hepatocytes, activated T-cells, skeletal muscle cells, macrophages, as well as cancer cell lines and immortalized cell lines. In our study, we find that *ATF3* expression is dependent on PenStrep and that it contains a PenStrep dependent regulatory region that may compound the regulatory effect of *ATF3* accessibility and expression. Given that *ATF3* expression may be significantly enhanced by PenStrep treatment, studies that examine *ATF3*’s role in pathways related to toxicity, cell proliferation and differentiation should avoid use of antibiotics in cell culture if possible.

The impact of culturing liver cells with antibiotics, as shown in our study on HepG2 cells, could also apply to other immortalized human cell lines that are commonly used to assess gene expression patterns at baseline and in drug-induced conditions and even primary cells. It is possible that antibiotics such as penicillin-streptomycin and gentamicin also induce a functional state that is significantly different from the basal state of these cell types. Further evaluation of the biological impact of antibiotic treatment across cell lines is highly warranted. However, we provide some evidence that using antibiotics in cell culture should be avoided-especially in studies focused on drug response as well as cell cycle regulation, differentiation, and growth. Data from studies in which antibiotics are used for cell culture should be examined with caution.

## Methods

### HepG2 cell culture

HepG2 cells (ATCC) were thawed at passage 37 and cultured in two separate batches with D-MEM, high glucose media (catalog # 11965-092, Life Technologies) that contained fetal bovine serum (catalog # CCFAP003, UCSF Cell Culture Facility) at 10%, L-glutamine (catalog # CCFGB002, UCSF Cell Culture Facility) at 1% in 6-well plates (catalog # 3516, Corning). One batch was cultured with this DMEM media supplemented with penicillin-streptomycin (catalog # CCFGK003, UCSF Cell Culture Facility) at the standard ATCC guideline of 1%. The other batch that served as the control group, was cultured in parallel in DMEM media that was not supplemented with penicillin-streptomycin and incubated at 37 degrees Celsius, 5% CO_2_ for two passages (or 21 days post-thaw).

### RNA-seq

HepG2 cells were cultured with PenStrep-supplemented media or control media in separate batches as mentioned above. The cells were then washed with PBS, and lysed directly with Buffer RLT from the RNAeasy mini kit (Qiagen) with the on-column DNase digestion step for collecting total RNA. Libraries were made with Illumina NGS library preparation for polyA tail selection on 5 ug of total RNA per sample by the UCSF Genomic Core Laboratories (http://humangenetics.ucsf.edu/genomics-services/). Three technical replicates were carried out for each condition. 50 bp single end sequencing was carried out on an Illumina Next-Generation Sequencing platform. The resulting reads were demultiplexed and aligned to the human genome (hg19) using STAR ^48^, which also calculated read counts for each gene in the Ensembl annotation. Analysis for differential expression across the nine replicates was performed using DESeq2 ^9^. DESeq2 was chosen due to its stringency in normalization and distribution methods. Gene ontology terms per RNA-seq cluster of DE genes were assessed using DAVID ^24^.

### Ingenuity pathway analysis

Ensembl Gene IDs and the log2FoldChange for the differentially expressed (DE) genes from our RNA-seq were uploaded into QIAGEN’s Ingenuity Pathway Analysis (IPA, QIAGEN Redwood City, http://www.ingenuity.com/). We used the following IPA core analysis settings: general settings, reference set = Ingenuity knowledge base (genes only), relationships to consider = direct and indirect relationships; data sources = all; confidence = experimentally observed only. We reviewed Ingenuity canonical pathways to study known biological pathways and processes of interest that were enriched in our DE genes. We also used the Ingenuity upstream analysis, which predicts the functional status of upstream regulators, such as transcription factors, kinases, and growth factors based on known downstream targets, i.e., the input set of the DE genes.

### ChIP-seq

Around twelve million cells were cultured separately from the RNA-seq experiments but in identical conditions as previously mentioned (37°C, 5% CO_2_ for two passages) on 6 well plates. For each immunoprecipitation, one plate was fixed with 1% formaldehyde for 15 minutes and quenched with 0.125 M glycine. The remainder of the ChIP-seq protocol was carried out using the Diagenode LowCell# ChIP kit (Diagenode; catalog number: C01010070) following the manufacturer’s protocol. Chromatin was isolated by adding lysis buffer and sheared to an average length of 300 bp with a Covaris sonicator. Genomic DNA regions of interest were isolated using 4 ug of antibody against H3K27ac (Abcam; catalog number ab4729). Genomic DNA complexes were washed, eluted from the beads with SDS buffer, and subjected to RNase and proteinase K treatment. Crosslinks were reversed by incubation at 65°C, and ChIP DNA was subsequently isolated and purified. Three technical replicates were carried out for each condition. Input genomic DNA was prepared by treating aliquots of chromatin with RNase, proteinase K and heat for de-crosslinking, followed by ethanol precipitation. Pellets were resuspended and the resulting DNA was quantified on a Bioanalyzer. ChIP and input DNAs were prepared for amplification using a ThruPLEX DNA-seq kit (Rubicon Genomics) following the manufacturer’s protocol. Library barcode adaptors were added to each sample during amplification and the library was size-selected (~150-200 bp) using a Diagenode iPure kit v2. The resulting amplified DNA was purified, quantified, and tested by a Bioanalyzer reading to assess the quality of the amplification reactions. Amplified DNA libraries were sequenced on and Illumina HiSeq 4000. All reads were mapped to the human genome using Bowtie ^49^. H3K27ac peaks were called against input using MACS ^50^ and peaks consistent across replicates identified using the ENCODE Irreproducibility Discovery Rate (IDR) pipeline ^51^. For differential peak intensity analysis, peaks across conditions were merged and reads coverage obtained using HTSeq ^52^. H3K27ac peaks differentially enriched in the PenStrep condition were then identified through DESeq2 ^9^ using a custom normalization matrix to correct for differing signal to noise ratios between replicates ^53^. As with the RNA-seq analysis, PenStrep dependent regions were identified through clustering analysis on all differentially enriched H3K27ac regions with adjusted p-values (FDR) <0.1 between the PenStrep-treated and non-treated conditions using the R package hclust and displayed in a heatmap.

### Cluster enrichment Analysis

To compare PenStrep dependent clusters of DE regions and DE genes, we used all 9,514 DE regions from the H3K27ac analysis and matched the closest gene expressed that had a normalized FPKM value >0 for all replicates across conditions. We cluster these regions into two groups (control/PenStrep dependent regions). Similarly, we cluster all DE genes (control/PenStrep responsive genes). We then test for Spearman’s correlation between H3K27ac signal in DE regions and RNA-seq signal in DE genes from these clusters and report the hypergeometric test enrichment p-values to determine the significance of overlap between PenStrep-dependent DE genes and regions.

## Acknowledgments

We thank Fumitaka Inoue, Navneet Matharu, and Nadja Makki for their troubleshooting advice on the ChIP-seq experiment, as well as John Rubenstein and Katherine Pollard for their feedback throughout this project. This work is supported in part by a grant from the National Institute of General Medical Sciences GM61390. NA is also supported by grants by the National Human Genome Research Institute and National Cancer Institute 1R01CA197139, National Institute of Mental Health 1R01MH109907 and National Institute of Child & Human Development 1P01HD084387. AHR was funded by the National Institute of General Medical Sciences Predoctoral Training Grant [T32 GM007175] and the National Science Foundation.

## Author information

**Affiliations**

**Department of Bioengineering and Therapeutic Sciences, University of California San Francisco, San Francisco, CA, USA**

Ann H. Ryu, Walter L. Eckalbar, Anat Kreimer & Nadav Ahituv

**Institute for Human Genetics, University of California San Francisco, San Francisco, CA, USA**

Ann H. Ryu, Walter L. Eckalbar & Nadav Ahituv

**Department of Electrical Engineering and Computer Science and the Center for Computational Biology, University of California Berkeley, Berkeley, CA, USA**

Anat Kreimer & Nir Yosef

## Contributions

AHR and NA conceived this project. AHR performed the experiments. AHR, WLE, AK, and NY performed the analysis. AHR and NA wrote the manuscript. All authors read and approved the final manuscript.

## Corresponding author

Correspondence to Nadav Ahituv.

